# MBASR: Workflow-simplified ancestral state reconstruction of discrete traits with MrBayes in the R environment

**DOI:** 10.1101/2021.01.10.426107

**Authors:** Steven Heritage

## Abstract

Ancestral state reconstruction (ASR) is a critical tool in comparative biology. Given a phylogenetic tree and observed trait states for taxa at the tree’s tips, ASR aims to model ancestral trait states at the tree’s internal nodes. Thus, the pattern, tempo, and frequency of a character’s evolution can be considered in a phylogenetic framework. Software programs that perform discrete trait ASR analyses are variably complex in terms of workflow steps, input formats, user interfaces, settings and execution commands, and the organization of output. ***MBASR*** (acronym for ***MrBayes Ancestral States with R***) is an R language toolkit that highly automates the ASR workflow and uses the machinery of the popular phylogenetics software MrBayes. Input data files are simple, analysis can be setup and run with a single R function, and statistical output is concisely organized and immediately interpretable. Use requires minimal knowledge of the most basic R commands and no experience with the MrBayes program. The ***MBASR*** toolkit can be freely downloaded from GitHub.

## Introduction

Several phylogenetics software programs include functionality to perform ancestral state reconstructions (ASR) of discrete trait data. The purpose of ASR is to estimate the state of a character in the past—in particular at ancestral nodes within a phylogenetic tree given character state observations at the tree’s tips and considering the tree’s topology and branch lengths. Discrete (qualitative) trait data might be binary (e.g., 0 or 1, denoting a character that is observed in two states; like absence or presence) or multistate (e.g., 0, 1, 2 or 3, denoting a character that is observed in four states; like red, yellow, green, or black). Additionally, some multistate characters might be conceived as “ordered” which can be a methodological approach to coding continuous (quantitative) traits by binning state observations into sequenced categories^1^ (e.g., 0, 1, or 2, denoting a size continuum; like small, medium, or large). When evaluated in a probabilistic architecture, discrete trait ASR returns likelihood values for each possible state of the character at internal nodes of the tree.

Software that performs discrete trait ASR can be standalone command-line or GUI programs like BayesTraits^2^, Mesquite^3^ and RASP^4^, or library packages like ape^5^ and phangorn^6^ for R^7^. These programs each have unique methods to load trait data and input trees, have their own commands, menus and settings, and write-out analysis results in different formats that are in various states of accessibility. Thus, even for experienced users, there is often a re-learning curve for the several steps needed to get data into an ASR analysis, getting it running, and wrangling the results into a useable and reportable packet.

***MBASR*** is an R toolkit that markedly simplifies this process. From within the R environment, ***MBASR*** performs ancestral state reconstruction using the MrBayes^8^ program and automates nearly all of the necessary steps. Calling a single R function will import the trait data and tree, execute the analysis, collect and format the results, and provide a tree plot with marginal likelihood pie charts at ancestral nodes. The toolkit leverages MrBayes’ robust implementation of the continuous-time Markov model^9,10^ for discrete trait evolution and also benefits from a large base of existing R users who are already comfortable with basic commands in the R language.

## Features

***MBASR*** is freely available from GitHub: https://github.com/stevenheritage/MBASR/releases. Package installation is not necessary. Two R dependency libraries (`ape` & `phytools`) are required and are automatically loaded (and installed, if absent). The toolkit comes with an existing folder structure that cannot be changed. The folder schema is self-evident—e.g., the folders “input.files” and “results” are where input data files are placed and analysis output is written. The user should place their executable file of MrBayes 3.2.7a (Windows or MacOS, serial version) in the folder named “mb”. The executable should be renamed “mb.exe” (Windows) or “mb” (MacOS). After launching R, the user prepares the environment by (1) assigning a path for the main ***MBASR*** folder and (2) loading a single source file. The toolkit is then ready for use.

Only two functions are intended for users (but several auxiliary functions are bundled and used by the algorithm). These functions and their arguments are:

### Function

~~~
MBASR(file.name.tree, file.name.trait.data,
file.name.plot.settings, character.type, n.samples)
~~~

### Description

This R function imports the trait data and tree, performs the ASR analysis, and writes-out two results files.

### Arguments

`file.name.tree` is the name of a tree file that resides in the “input.files” folder. The tree must be rooted and include branch lengths. The Newick tree format is required. If needed, a tree can be prepared (root, re-root, convert to Newick) using the software FigTree^11^. In FigTree, click on any branch desired and then on re-root. Then select: File > Export Trees > Tree File Format : Newick > Save As Currently Displayed > OK.

`file.name.trait.data` is the name of a trait data file that resides in the “input.files” folder. Trait scores are contained in a simple text file. Each line in the file contains a single taxon name, followed by a tab, followed by a state score (e.g., 0 or 1 or 2). The taxa in the trait data file can be in any sequence but all taxa that are in the tree must also be present in the trait data file. Missing scores are allowed and should be coded as a question mark (?). Ambiguity scores are also allowed and should be coded using an ampersand (e.g., 1&2). Trait scores must follow standard coding—use 0 through 9 for unordered characters; use 0 through 5 for ordered characters. Taxa names in the trait data file and tree file must be identical. Avoid using spaces, periods, or quotes in names.

`file.name.plot.settings` is the name of a simple text file that resides in the “input.files” folder. Example versions of this file are provided and can be copied and edited as desired. The values in the plot settings file define cosmetic attributes for the tree plot like font size for taxa names and the size of pie charts at tree nodes. Default settings are suggested, but these might not be an appropriate choice for trees of all sizes.

`character.type` can be set to “unordered” or “ordered”, depending on the nature of the character being evaluated.

`n.samples` is an integer value that designates how many samples will be generated. The MrBayes MCMC mechanism has been pre-configured to sample every 100 generations (i.e., if `n.samples=`500 then MrBayes ngens will be 50000). Very large samples are probably unnecessary—1000, 10000 and 100000 samples can yield nearly identical results.

### Function

~~~
replot(file.name.tree, file.name.trait.data,
file.name.plot.settings)
~~~

### Description

This R function replots the tree after the user has run an ASR analysis and subsequently edited/adjusted the plot settings file (in the “input.files” folder). In other words, a tree plot’s cosmetic attributes (e.g., font size, pie chart size, etc.) can be changed without re-running the ASR analysis. To do so, be certain ***not*** to delete, move, or rename the file “MrBayes.ASR.results.txt” which was written to the “results” folder.

### Arguments

`file.name.tree` is the same as above.

`file.name.trait.data` is the same as above.

`file.name.plot.settings` is the same as above.

***MBASR*** can be used to reconstruct one trait per run. Output of the `MBASR` function is written to the “results” folder. The output “MrBayes.ASR.results.txt” is a tab delimited text file that includes node numbers in the first column and probabilities of states in subsequent columns. If a character is coded with three states (0, 1 and 2) then the output will contain three columns to the right of the node numbers column. Node numbers correspond to the input tree and follow the standard conventions used by BayesTraits and all R phylogenetics packages (e.g., ape, phangorn, phytools^12^, and paleotree^13^). If there are 100 tips on a tree, the root split will be node number 101—this will be the first node number in the results table. State probabilities are calculated as the means of post-filtered generations from the MrBayes MCMC sampler. Probabilities across any row in the results table will sum to 1. The output “tree.plot.pdf” is a standard PDF file with vector art that can be opened with several viewers and editors. If the intent is to open the PDF with Adobe Illustrator, be sure to install the open type font “AdobePiStd” to avoid font substitutions and flawed rendering.

The ***MBASR*** algorithm uses a single input tree to hard-constrain the topology and node ages during the ASR analysis. With no uncertainty in either of these parameters, the MrBayes proposals to re-estimate *Tau* and *V* (topology and branch lengths) are disabled for the call. MrBayes’ native Markov model is used for the analysis with variable coding enabled to accommodate ascertainment bias^9,14^. The evolution of only a single character is modeled within a fixed phylogeny, so very large samples are probably unnecessary. Given these pre-configured analysis parameters, the statistical results will more closely resemble a maximum likelihood estimate of state marginal likelihoods than a Bayesian estimate from a posterior distribution (containing variation in tree topology and branch lengths) that is summarized with HPD ranges of uncertainty. An automatic filter discards burnin samples (and poor estimates) by identifying the range of generation log likelihoods in the MCMC posterior set and then removing generations that score below the upper 25% threshold. Changing the character type from “unordered” to “ordered” can substantially impact the calculation of likelihoods when the trait transforms to non-adjacent states; doing so can considerably change the ASR results.

## Conclusion

Ancestral state reconstruction analysis is widely used in phylogenetic comparative research. Typical applications include investigating the patterns of a trait’s evolution using an existing tree, identifying morphological character support after a phylogenetic analysis has been run, and modeling the geographic origins of clades within a taxonomic group of interest^15^ (Fig. 1). The MrBayes mechanism for discrete trait ASR uses continuous-time Markov modeling against a tree’s topology and branch lengths to provide a statistical estimate for character states at ancestral tree nodes. This approach differs significantly from non-statistical character optimization and other methods that do not include a tree’s branch lengths in calculations. The ***MBASR*** toolkit wraps MrBayes’ ASR functionality while streamlining and accelerating a user’s workflow by using the simplest input data formats, calling a single function for setup and execution of the analysis, and automating the collection and summarization of results into concise and directly useable output.

**Figure 1.**
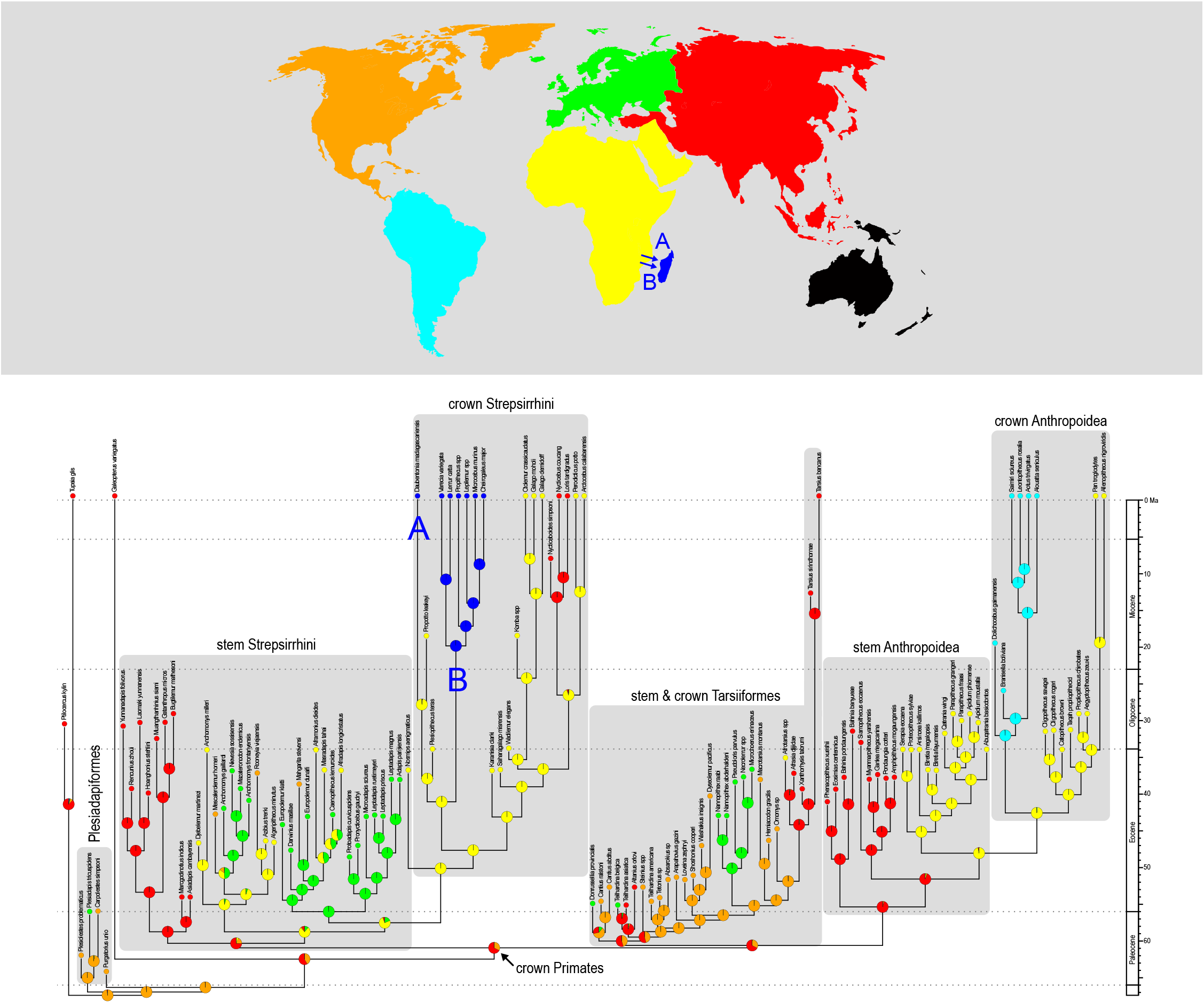
MBASR ancestral state reconstruction of geographic landmass for living and fossil primates. Tree and trait scores are from Gunnell et al. (2018) and are bundled with the MBASR toolkit as example data. The MBASR algorithm is very similar to the method and settings used in that study. The toolkit outputs a simple text table that reports the probabilities of each trait state at each ancestral tree node. Output also includes a vector art tree plot that overlays probability pie charts at lineage splits. In this figure, additional annotations have been added (e.g., map & timescale).

## Acknowledgements

Erik Seiffert and Matthew Borths beta-tested the functions and provided useful feedback. Emmanuel Paradis and several co-authors wrote the dependency package *ape*. Liam Revell wrote the dependency package *phytools*. John Huelsenbeck, Fredrik Ronquist, Paul van der Mark, Maxim Teslenko, Chi Zhang and several other co-authors wrote the *MrBayes* software.

## Additional Information and Declarations

### Availability

The *MBASR* toolkit can be freely downloaded at GitHub: https://github.com/stevenheritage/MBASR/releases The requisite *MrBayes* software is linked here: http://nbisweden.github.io/MrBayes/download.html *R* is available at: https://www.r-project.org/

